# Shift-inducible [transgenerational] increase in recombination rate as an evolving strategy in a periodic environment: a numerical model

**DOI:** 10.1101/2021.10.28.466217

**Authors:** Sviatoslav Rybnikov, Sariel Hübner, Abraham B. Korol

## Abstract

Numerous empirical studies have witnessed a plastic increase in meiotic recombination rate in organisms experiencing physiological stress due to unfavourable environmental conditions. Yet, it is not clear enough which characteristics of an ecological factor (intensity, duration, variability, etc.) make it stressogenic and therefore recombinogenic for an organism. Several previous theoretical models proceeded from the assumption that organisms increase their recombination rate when the environment becomes more severe, and demonstrated the evolutionary advantage of such recombination strategy. Here we explore another stress-associated recombination strategy, implying a reversible increase in recombination rate each time when the environment alternates. We allow such plastic changes in the organisms, grown in an environment different from that of their parents, and, optionally, also in their offspring. We show that such shift-inducible recombination is always favoured over intermediate constant optimal recombination. Besides, it sometimes outcompetes also zero and free optimal constant recombination, therefore making selection on recombination less polarized. Shift-inducible strategies with a longer, transgenerational plastic effect, are favoured under slightly stronger selection and longer period. These results hold for both panmixia and partial selfing, although selfing makes the dynamics of recombination modifier alleles faster. Our results suggest that epigenetic factors, presumably underlying the environmental plasticity of recombination, may play an important evolutionary role.

## 1. Introduction

The concept of environmental plasticity refers to the ability of a genotype to exhibit different phenotypes when exposed to different environments. The meiotic recombination rate (RR) has been recognized as a plastic trait for more than a century since Plough obtained a clear-cut U-shaped curve by plotting *Drosophila* RR in a certain genome region against the cultivation temperature [1]. Later experiments showed that, even if an environmental factor is handled with a coarse resolution, insufficient to figure out the whole continuous reaction norm, the pattern is typically consistent: RR tends to increase when the organism is stressed. To date, such stress-induced increase in RR is documented for various species (reviewed. in [2–4]). The so far proven recombinogenic factors include not only abiotic stressors like cold, heat, starvation, and desiccation, but also parasite infection [5,6], and even behavioural stressors like overpopulation [7].

The recombinogenic effect of ‘primary’, physical stressors may arise from changes in biochemical pathways in the germline cells (reviewed in [8,9]). ‘Higher-level’, behavioural stressors, probably also involve complex soma-to-germline signalling, whose existence attaches ever-increasing attention, especially in the context of epigenetic variation [10–13]. Yet, regardless of the mechanistic explanations, an intriguing question is whether stress-inducible recombination could have evolved as an adaptive trait. To address it, here we model a diploid population in a two-state periodic environment, with each environmental shift causing physiological stress in all population members. In this setup, we trace the dynamics of selectively neutral modifier genes conferring shift-inducible recombination. Several theoretical studies earlier demonstrated the evolutionary advantage of plastic recombination in a periodic environment [14–16]. However, they assumed rather complex dependencies of RR on environmental stressors, allowing benefits not only from the shift-inducible increase but also from additional mechanisms. For example, the stress-associated increase in RR was genotype-specific [15,16] or lasted during all generations of one of the environmental states (defined as ‘stressful’) rather than followed the environmental shifts [14,16]. Now, we aim to dissect the advantage of shift-inducible recombination per se. To maximize consistency with our previous study [16], we consider a similar three-locus selected system and a similar two-state environment. However, we now exclude the environmental asymmetry: the two states differ only in the direction but not the intensity of selection so that none of them can be defined as stressful. We also exclude benefits from the between-genotype variation: after each environmental shift, the RR increases equally in all organisms.

Recent reviews agree that recombination plasticity has already become a commonly recognized phenomenon and suggest switching the research focus from its existence to its quantitative characteristics; a key task within the new agenda should be revealing the evolutionary forces affecting the persistence and the strength of plastic changes in RR [17,18]. Sharing this viewpoint, here we examine shift-inducible recombination with different duration of the plastic effect, including transgenerational one. In his pioneering study on the consequences of larval starvation in *Drosophila*, Neel [19] observed a persistent, life-long increase in RR in the enclosed heterozygous females, suggesting that effects of the stressor somehow passed through many cell cycles in their germline. Such a lag between the stressor exposure and the recombination response was soon documented also for high temperature, in fruit flies [20,21] and fungi [22–24]. Moreover, later studies reported an increased RR even in organisms who were never exposed to a stressor themselves but originated from the stressed parents [25,26]. Recent experiments suggest that even epigenetic markers associated with increased somatic recombination are capable to pass through some rounds of meiosis, causing an effect during several consequent generations [27,28]. Although empirical evidence for this phenomenon remains extremely limited, we found it highly perspective to explore the evolvability of recombination’s transgenerational plasticity theoretically, to provoke further empiric investigations. Thus, here we also test for the evolutionary advantage of shift-inducible recombination with a longer plastic increase in RR, lasting two generations after the environmental shift, either with or without damping.

We conduct all simulations for populations with panmixia and partial selfing, given the earlier revealed modulating effect of the mating system on the evolution of recombination [29–31], including plastic recombination [32].

## 2. Model

### (a) Life cycle

We consider an infinite population of obligatory sexual diploids. The generations do not overlap. Each generation includes (*i*) maturation from zygotes to adults, when viability selection acts, (*ii*) gametogenesis, when recombination occurs, and (*iii*) reproduction via one of the considered mating strategies. Each mating strategy is defined by the fraction α of adults practicing selfing (the rate of selfing); the rest 1-*α* adults practice outcrossing. We examined three mating strategies: panmixia (*α* = 0) and partial selfing with *α* = 0.25 and *α* = 0.5 (compete selfing was not considered since in the absence of mutations it quickly eliminates heterozygotes, leaving no place for the evolution of recombination). The rate of selfing was treated as a parameter – in contrast to the rate of recombination treated as a function of the genotype at a specific modifier locus (see below).

### (b) Selected system and selection regime

Each organism bears three bi-allelic loci: *A*_1_/*A*_2_, *B*_1_/*B*_2_, and *C*_1_/*C*_2_, that together encode a quantitative trait *Q*. At each locus *L* (*L* = {*A, B, C*}), the homozygotes *L*_1_*L*_1_ and *L*_2_*L*_2_ display locus-associated trait values *q_L_* = 0 and *q_L_* = 1, respectively. The three loci contribute to the trait in a purely additive way: *Q* = *∑_L_ q_L_*. The overall trait value varies, therefore, from *Q*_min_ =0 (in the genotype *A*_1_*A*_1_*B*_1_*B*_1_*C*_1_*C*_1_) to *Q*_max_ =3 (in the genotype *A*_2_*A*_2_*B*_2_*B*_2_*C*_2_*C*_2_).

The environment has two alternating states and fluctuates with period 2*T*. The two states are equal in the duration (each lasts *T* generations) and the selection intensity *s* but are opposite in the optimal trait value *Q*_Opt_ (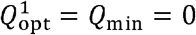 in the state 1 and 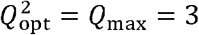 in the state 2). In both states, genotype absolute fitness *W* declines in a Gaussian-like way with the trait *Q* departing from the optimum *Q*_Opt_: *W*(*Q*) = exp{−*Δ*Q^2^/2*σ*^2^}. The fittest genotypes (*A*_1_*A*_1_*B*_1_*B*_1_*C*_1_*C*_1_ in state 1 and *A*_2_*A*_2_*B*_2_*B*_2_*C*_2_*C*_2_ in state 2) have absolute fitness *W*_max_ = 1, while the least fit genotypes (those with Δ*Q* = Δ*Q*_max_ = 3) have absolute fitness *W*_min_. The selection intensity s is defined in an intuitively clear way, as s=1 − *W*_min_. If required, the parameter *σ* in the above-mentioned expression for fitness can be expressed as 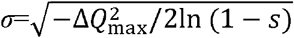.

The polymorphism within the selected system is maintained by the temporally varying selection. Yet, to support it further (which appeared necessary under high selfing – see Section 3*d*), we exploited dominance lift [33], putting equal *d* = 0.25. Thus, the heterozygote *L*_1_*L*_2_ display locus-associated trait value 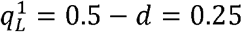 in state 1 (when allele *L*_1_ is favored) but 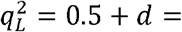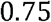 in state 2 (when allele *L*_2_ is favored) – instead of 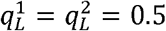 in both states.

We examined selection regimes with half-periods *T* = 2, …,15 generations and selection intensity *s* = 0.01, …, 1 with the step of 0.01 (as a proxy for *s* = 1 we used *s* = 0.999, since the Gaussian function is not defined for *s* = 1).

### (c) Recombination strategies

The recombination strategy of each organism is defined by the genotype at a specific selectively neutral locus (‘recombination modifier’), either unlined or linked to the selected system. Each strategy is described by RRs in the first *r*_1_, second *r*_2_, and all consequent *r*_3+_ generations after the environmental shifts. We considered two types of strategies: *constant* recombination, with the same rate over the period (*r*_1_ = *r*_2_ = *r*_3+_ = *r*), and *shift-inducible* recombination, with the rate increased in at least one generation after the environmental shifts (*r*_1_ = *r* + Δ*r*). In the latter case, the plastic increase may be limited to *one generation* (*r*_1_ = *r* + Δ*r*, *r*_2+_ = *r*) or be *transgenerational*, either damped, with twice weaker plastic changes in the second generation after the shift (*r*_1_ = *r* + Δ*r*, *r*_2_ = *r* + Δ*r*/2, *r*_3+_ = *r*) or non-damped, with similar plastic changes during two generations after the shift (*r*_1_ = *r*_2_ = *r* + Δ*r*, *r*_3+_ = *r*). The magnitude of the plastic increase Δ*r* = 0.05.

All recombination strategies were compared based on the dynamics of the corresponding modifier alleles [34]. One strategy was considered to be favoured over another if its allele both invaded the population from rarity (0.05) and resisted the backward invasion. In such pairwise comparisons, the modifier allele dynamics were traced during 2,000 periods following 2,000 periods of the evolution with the monomorphic modifier. The modifier allele in question was always injected in Hardy-Weinberg equilibrium with the opponent allele and linkage equilibrium with the selected genotypes.

We first estimated the optimal constant recombination and then tested whether shift-inducible recombination can be even a better strategy. Shift-inducible recombination always implied a higher RR after the environmental shift, compared to that in the rest part of the period. At the same time, shift-inducible recombination could be characterized by either lower or higher mean RR than the corresponding competing constant recombination. Hereafter we refer to two forms of shift-inducible recombination as inferior and superior strategies respectively. Zero optimal constant recombination was compared with the superior strategy, implying the increase by Δ*r* = 0.05 after the environmental shifts and further return to zero recombination. Free optimal constant recombination was compared with the inferior strategy, implying free recombination after the environmental shifts and the further decrease by Δ*r* = 0.05. Intermediate optimal constant recombination was compared with both inferior and superior strategies (Fig. 1).

**Figure 1.**
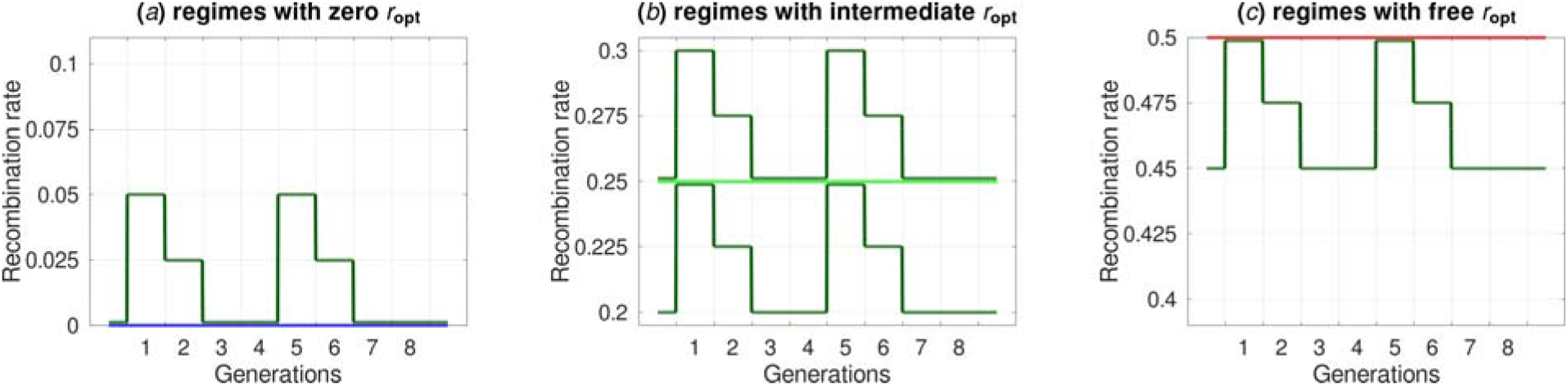
The examined pairs of competing recombination strategies: *(a)* Zero optimal constant recombination (blue line) vs. superior shift-inducible recombination (dark green line); *(b)* Intermediate optimal constant recombination (light green line) vs. inferior and superior shift-inducible recombination (lower and upper dark green lines, respectively); *(c)* Free optimal constant recombination (red line) vs. inferior shift-inducible recombination (dark green line). The example shows an environment with half-period *T* = 4 generations and the damped trans-generational shift-inducible strategy. In the case of the one-generation strategy, the increase in recombination rate in the second generation after the environmental shift is absent, while in the case of the non-damped transgenerational strategy, the increase persists for two generations after the shift.

## 3. Results

### (a) Panmixia: constant recombination

For each selection regime, we first compared different constant strategies to evaluate the optimal constant RR (*r*_opt_). In the prevailing proportion of regimes, with a relatively short period or weak selection, any constant recombination was rejected (*r*_opt_ = 0). Less frequently but still quite often, under a relatively long period and strong selection, free recombination was favoured (*r*_opt_ = 0.5). Thus, the two parameters, selection intensity *s*, and period duration *T*, affected the optimal constant RR *r*_opt_ unidirectionally, suggesting that their product s*T* can efficiently predict the direction of selection on recombination; hereafter, we refer to this product as ‘cumulative selection’.

Between the two above areas (with *r*_opt_ = 0 and *r*_opt_ = 0.5), a narrow strip of intermediate optimal constant recombination (0 < *r*_opt_ < 0.5) occurred. Its width depended on the modifier linkage to the selected system, being the highest for the intermediately linked (*r*_M_ = 0.2) and absent for the unlinked (*r*_M_ = 0.5) modifier. Sometimes, under strong selection but short period, we observed bistability – a situation when either *r*_opt_ = 0 or *r*_opt_ = 0.5 behaved as a local optimum, depending on the initial RR in the population. Figure 2*a* summarizes the effect of modifier linkage *r*_M_ on the distribution of selection regimes by the optimal constant RR *r*_opt_, while Figures 2*b* and 2*c* show the effects of period and selection intensity on *r*_opt_ for two chosen values of the modifier linkage: *r*_M_ = 0.2 and *r*_M_ = 0.5.

**Figure 2.**
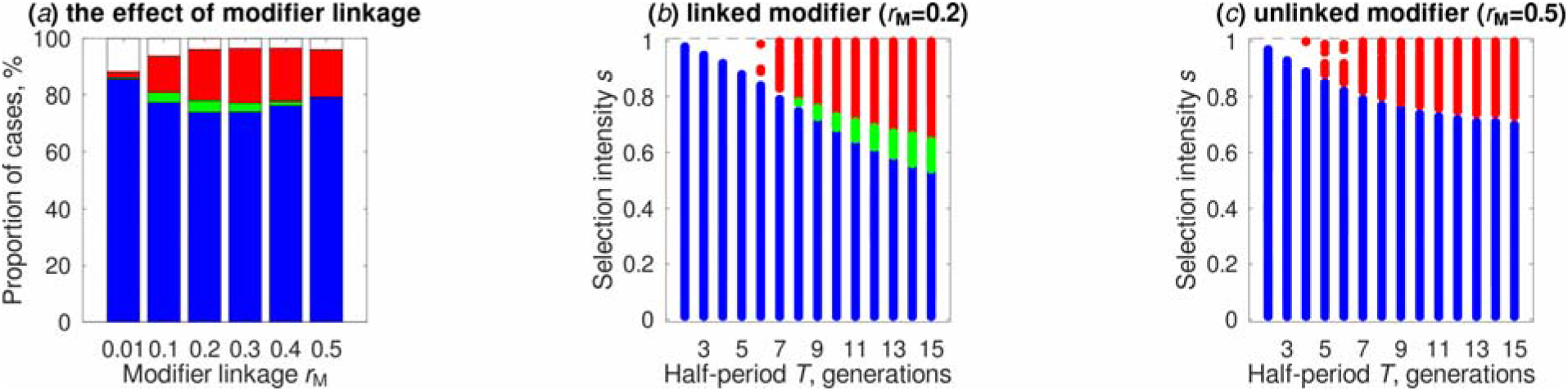
The effect of modifier linkage on the distribution of selection regimes by the optimal constant recombination rate (a), and the effects of period and selection intensity on this rate under linked (*b*) and unlinked (c) modifier. The colours stand for zero (blue), intermediate (light green), and free (red) optimal constant recombination; selection regimes with bistability (i.e., those where either zero or free recombination appeared as a local optimum, depending on the initial recombination rate in the population) are left uncoloured.

### (b) Panmixia: one-generation shift-inducible recombination

The found optimal constant recombination was compared with shift-inducible recombination to test whether the latter is even more advantageous. We first tested shift-inducible recombination with one-generation plastic increase (i.e., in the organisms, grown in an environment different from that of their parents). Such strategies always outcompeted the intermediate optimal constant recombination, whenever the latter existed (Fig. 3*b*). The advantage over zero optimal constant recombination was observed only under certain conditions, namely a relatively long period and strong selection (among those ensuring *r*_opt_ = 0) (Fig. 3*a*). In contrast, the advantage over free optimal constant recombination occurred under a relatively short period and weak selection (among those ensuring *r*_opt_ = 0.5) (Fig. 3*c*). Yet, in both these comparisons (against *r*_opt_ = 0 and against *r*_opt_ = 0.5), shift-inducible recombination outcompeted in the parameter areas adjacent to the area of intermediate optimal constant recombination. Since these plastic strategies made the population’s mean RR intermediate (moved it either upward from *r* = 0 or downward *r* = 0.5), this extended the area of intermediate optimal recombination, thereby making the selection for/against recombination less polarized.

**Figure 3.**
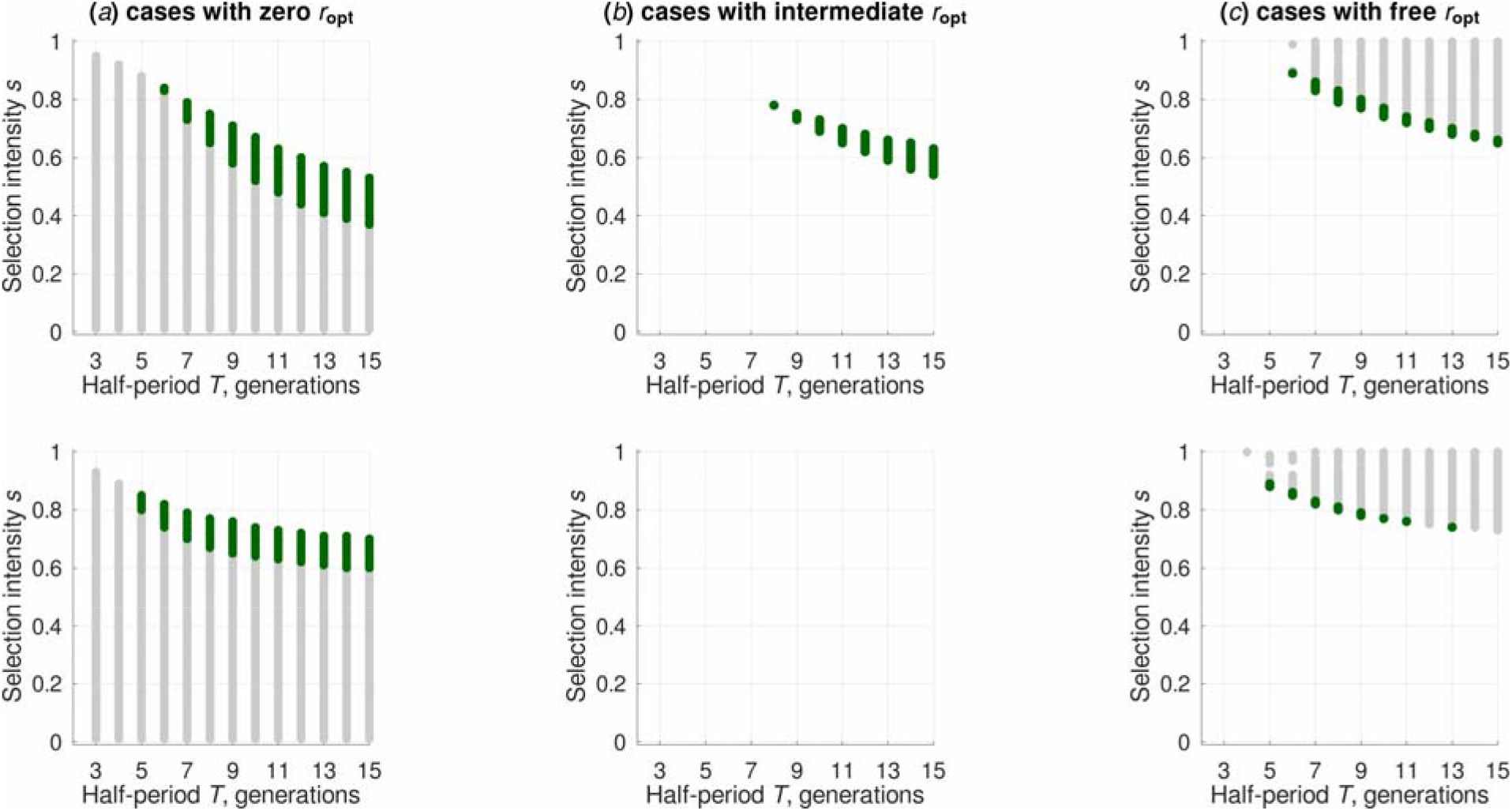
The evolutionary advantage of shift-inducible recombination (green points) over zero (*a*), intermediate (*b*), and free (*c*) optimal constant recombination, for linked (upper row) and unlinked (bottom row) modifier. Selection regimes with intermediate optimal constant recombination were found only with linked modifier (see Fig. 2(b) & 2(c)).

### (c) Panmixia: transgenerational shift-inducible recombination

We then examined transgenerational shift-inducible recombination, with a plastic increase in RR during two generations after the environmental shift (i.e., not only in the organisms, grown in an environment different from that of their parents but also in their offspring). The transgenerational strategies always demonstrated a sounder advantage over the corresponding intermediate optimal constant recombination than the one-generation strategies; moreover, the transgenerational strategies outcompeted the one-generation strategy in the direct comparisons. However, in competition with zero and free optimal constant recombination, the advantage of transgenerational strategies depended on selection parameters. When recombination was selected against (*r*_opt_ = 0), transgenerational strategies outcompeted in a slightly narrower parameter area than those with a one-generation increase (Table 1, the upper part). The reason is that too short period and too weak selection made too long plastic increase unnecessary and even disadvantageous, while a shorter increase still remained favoured. In contrast, when recombination was selected for (*r*_opt_ = 0.5), strategies with a longer plastic increase better supplied the evolutionary demand in recombination. In this area, transgenerational strategies appeared to be favoured more frequently than those with a one-generation increase (Table 1, the bottom part). On average, shift-inducible strategies with longer plastic effect were favoured under stronger cumulative selection 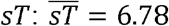 for one-generation strategy, 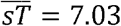 for damped transgenerational strategy, and 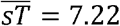 for non-damped transgenerational strategy. For the unlinked modifier, the corresponding numbers were 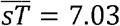, 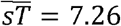, and 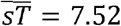.

**Table 1.** The proportion of selection regimes (%) where shift-inducible recombination was favoured over the optimal constant recombination*

| Shift-inducible strategy | Panmixia |  | 25% selfing |  | 50% selfing** |  |
| --- | --- | --- | --- | --- | --- | --- |
|  | Linked modifier | Unlinked modifier | Linked modifier | Unlinked modifier | Linked modifier | Unlinked modifier |
| Among selection regimes with *zero* optimal constant recombination |  |  |  |  |  |  |
| One-generation strategy | 14.3 | 11.4 | 15.2 | 19.4 | 12.2 | 17.6 |
| Damped transgenerational strategy | 12.2 | 10.2 | 12.9 | 16.9 | 10.1 | 15.4 |
| Non-damped transgenerational strategy | 11.4 | 9.4 | 12.0 | 16.3 | 9.1 | 14.4 |
| Among selection regimes with *intermediate* optimal constant recombination |  |  |  |  |  |  |
| One-generation strategy | 100.0 | – | 100.0 | 100.0 | 100.0 | 100.0 |
| Damped transgenerational strategy | 100.0 | – | 100.0 | 100.0 | 100.0 | 100.0 |
| Non-damped transgenerational strategy | 100.0 | – | 100.0 | 100.0 | 100.0 | 100.0 |
| Among selection regimes with *free* optimal constant recombination |  |  |  |  |  |  |
| One-generation strategy | 12.3 | 5.5 | 11.1 | 6.2 | 9.4 | 10.0 |
| Damped transgenerational strategy | 19.4 | 10.6 | 17.8 | 12.8 | 15.5 | 15.1 |
| Non-damped transgenerational strategy | 27.3 | 18.2 | 25.6 | 19.0 | 23.1 | 21.5 |

\* Excluding selection regimes with $T = 2$ , where shift-inducible recombination with one-generation but not transgenerational plastic increase could be considered.
\*\* Including selection regimes where $r_{\text{opt}} = 0$ appeared as local rather than the global optimum.

Remarkably, the parameter areas where shift-inducible recombination was favoured over zero and free optimal constant RR, appeared to be adjacent to the area of intermediate optimal constant RR. Thus, recombination plasticity extended the area of intermediate constant recombination, making selection on recombination less ‘polarized’. For example, intermediate constant recombination was observed only in less than 4% of the examined selection regimes with the linked modifier, and was absent with the unlinked modifier (Fig. 2(*b*) and 2(*c*)); with shift-inducible recombination, the area of intermediate mean RR exceeded 17% and 10%, respectively.

### (d) Deviation from panmixia

We finally examined to which extent the above results are robust to deviations from panmixia. It turned out that selfing embarrasses polymorphism maintenance within the selected system. Starting from a certain rate of selfing, too high recombination led to allele fixation in two of the three selected loci, and this polymorphism-ruining RR declined with selfing. At the same time, dominance lift served as a counterwork, making higher selfing and higher recombination compatible with polymorphism. The herein used dominance lift *d* = 0.25 ensured recombination-independent polymorphism for selfing *α* ≤ 0.45. For selfing *α* = 0.5, a minor proportion of selection regimes lost polymorphism under very high recombination; yet this did not crucially affect the results since the optimal constant recombination in such regimes was zero.

Overall, the results for partial selfing appeared qualitatively similar to those for panmixia (Table 1). Notably, in selection regimes with *r*_opt_ = 0, shift-inducible recombination was favoured most frequently under intermediate selfing, regardless of the duration of the plastic effect and modifier linkage. At the same time, selfing exerted a sound effect on the speed of modifier dynamics, facilitating the invasion of the allele for shift-inducible recombination (Fig. 4).

**Figure 4.**
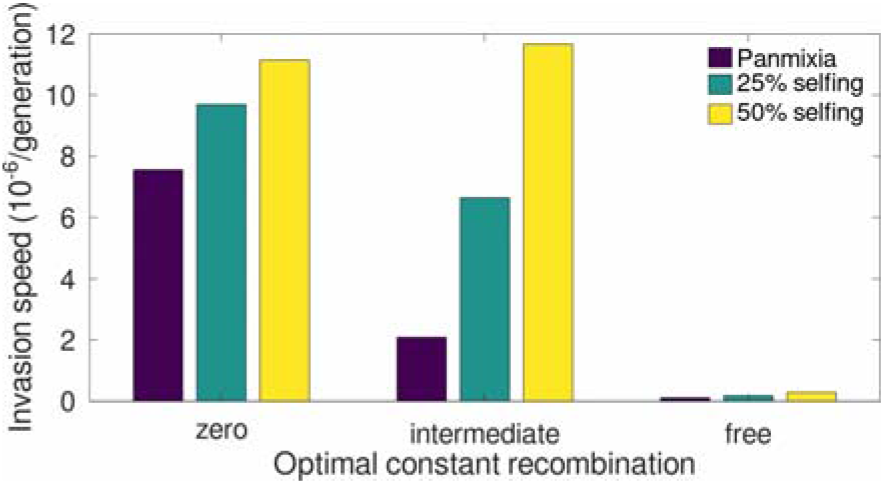
The effect of selfing on mean invasion speed of the favourable modifier allele for shift-inducible recombination. The data stand for non-damped transgenerational strategy; the modifier is linked to the selected system.

## 4. Discussion

### (a) Shift-inducible recombination in the context of stress-induced variation

In this study, we tested for the evolutionary advantage of shift-inducible recombination, implying an increase in RR after each environmental shift. We view such a recombination strategy in a broader context of stress-associated variation. Although stress is widely recognised as a potent evolutionary force [35–37], the interplay between different aspects of stress-induced variation is poorly explored. Two previous studies examine the evolution of stress-associated recombination in a two-state asymmetric periodic environment, with one state (either atypical [14] or that with strong selection [16]) defined as stressful. Therein considered recombination strategy implied an increase in RR during the whole stressful state, as compared to the other, benign state. However, such a formalization neglects the fact that the organisms may adapt to the environment physiologically and therefore experience milder stress at the end of the stressful period compared to its beginning. The fitness-dependent recombination, with the RR proportional to individual fitness [15,16], is free of the above flaw. However, such recombination strategy implies, in addition to the between-environment component, also the between-genotype component of RR variation, which embarrasses inferring on the primary sources of the observed benefits. A pure setup should allow for only one component of RR variation. Thus, to demonstrate the advantage of the between-genotype variation in RR alone, one should assume a constant environment, which was indeed done in several mutation-selection balance models [38,39]. In the current study, we, in contrast, exclude the between-genotype variation in RR: The only source of stress in our model is environmental shifts, and this transient stress equally affects all population members. The above-presented simulation results demonstrate that such shift-inducible recombination appears an evolutionarily advantageous strategy under a wide range of examined parameters.

### (b) Mechanisms behind the evolutionary advantage of shift-inducible recombination

In our simulations, shift-inducible recombination was always favoured over intermediate optimal constant RR. Importantly, this holds for both inferior and superior strategies of shift-inducible recombination. Such a successful pass of this so-called ‘two-reference’ test offered by Agrawal *et al.* [40] ensures that the observed advantage comes from RR plasticity *per se*, rather than from accidental changes in the population’s mean RR that might emerge as an uncontrolled by-product.

Mechanistically, a recombination modifier allele can spread in the population owing to either tightening its linkage to the favourable selected system (direct effect) or creating more favourable selected systems (indirect effect) [2,40,41]. This holds for modifiers of both constant and plastic recombination. At that, to evolve, two arbitrary recombination modifier alleles must confer different RRs in the selected system they ‘serve’ (the necessity of indirect effect); otherwise, they will remain neutral to one another, even if they confer different RR between themselves and the selected system (the insufficiency of direct effect) [16]. However, when the indirect effect is present, the direct effect stops being neutral and may also contribute to the recombination modifier dynamics. Such an interplay between the two effects is especially important for the evolution of plastic recombination. In this case, the varying RR within the selected system implicitly causes variation in RR between the modifier alleles and non-adjacent selected loci, even though the modifier’s linkage to the adjacent selected locus (i.e., direct effect in its narrow sense) is not explicitly assumed [39]. Bearing in mind the above regularities, in our simulations with panmixia and unlinked modifier, the observed advantage of shift-inducible recombination should be assigned to the plastic indirect effect alone. At the same time, in the simulations with the linked modifier, the advantage came from both the plastic indirect effect and the implicit, ‘plasticity-induced’ direct effect. Probably, these two effects jointly worked also in the simulations with partial selfing, given that the latter acts in a certain sense similar to the linkage, strengthening the modifier’s association with the selected system [31,42].

The advantage of shift-inducible recombination over zero and free optimal constant RR was observed only under certain parameter combinations and required respectively relatively strong and relatively weak cumulative selection (expressed as the product s*T*). The reason is that in both these situations, shift-inducible recombination moved the population’s mean RR from the optimal value (either upwards from *r*_opt_ = 0 or downwards from *r*_opt_ = 0.5). Thus, to be favoured, shift-inducible recombination had to exhibit stronger benefits from RR plasticity compared to these (‘non-optimality’) costs. It seems plausible to expect these costs to be lower under weaker selection for mean RR, which happens under relatively high values of sT in case of *r*_opt_ = 0, or relatively low values of s*T* in case of *r*_opt_ = 0.5. We also tried to link these findings with two important population-level characteristics, often targeted in models with the periodic environment in general and those focusing on the evolution of recombination in particular: Geometric mean of the population’s mean fitness across the period and range of the population’s mean fitness within the period [43,44]. It turned out that the range of fitness is the most informative predictor for the fate (advantage *vs.* disadvantage) of shift-inducible recombination, while geometric mean appeared negligibly informative (the relative explanatory effects ~95% and ~5% of the explained variation, respectively – not shown). Indeed, with stronger selection and longer period (higher values of s*T*), a larger proportion of the population is represented by favourable selected genotypes at the end of each environmental state. This ensures high mean fitness before the environmental shifts and therefore low fitness after them when the selection direction opposes. These two values of mean fitness (before and after the shifts) are respectively the highest and the lowest within the period, and thus define the range. Apparently, a larger range creates a stronger demand in recombination to ‘recover’ the favourable population structure. The central role of the fitness range is remarkably in line with the result of Charlesworth [45], even though the environment in his model changed continuously, not discretely.

In our simulations, transgenerational shift-inducible strategies appeared to be favoured under slightly stronger cumulative selection (that is, under longer period and stronger selection in each generation) than one-generation strategies; moreover, non-damped transgenerational strategies required stronger selection than the damped ones. This finding remarkably concurs with the conclusion by Furrow and Feldman [46] that longer environmental fluctuations facilitate selection for more persistent epigenetic regulation. Moreover, our results point to the possibility to extend those by Furrow and Feldman to the domain of selectively neutral variation. In this sense, our results corroborate those of Lobinska et al. [47] who have recently theoretically demonstrated the evolvability of epigenetically inherited mutations, as a strategy for maximizing the rate of adaptation.

## 5. Biological relevance of the model and conclusions

Molecular mechanisms enabling recombination plasticity are extremely sophisticated and underexplored, let alone those ensuring its modulation and evolution. The recombination control system includes recombination enzymes themselves (the acting subsystem) and chromatin organization in meiotic cells upon which the enzymes work (the reacting subsystem), and both these subsystems are subject to a joint genetic and epigenetic regulation. Several genome-wide association studies have revealed numerous candidate genes controlling meiotic recombination, including cohesion, axis, and synaptonemal-complex proteins [48–51]. Among them, a key modulator of recombination plasticity is probably NEC8, given its wide functionality (including an effect on the local chromatin structure) and sensitivity to oxidative stress [52,53]. The chromatin-based regulation is enabled by reversible epigenetic marking, such as methylation of cytosine and histones and acetylation of histones, which modulates the chromatin compaction and accessibility to enzymes involved in DNA metabolism, including recombination enzymes (reviewed in [54,55]). Several studies argue for a negative methylation-recombination association so that elevated RRs collocate with hypomethylated genome regions and vice versa [56,57]. In turn, the methylation status is likely fine-tuned by small interfering RNAs [58–60]. These small regulatory molecules are probably also triggered by the oxidative burst in the stressed cells [61]. They are capable of migrating into the neighboring untreated tissues potentiating a systemic organism-wide stress response. Moreover, they can migrate to the germline cells [62,63] and further to gametes and the next-generation zygotes [64], thereby ensuring a transgenerational increase in RR. Additionally, recombination plasticity can be affected by some ‘third-party’ competing processes. For example, RRs tend to negatively correlate with the activity of transposable elements [65], while the latter is also affected by environmental stressors via epigenetic regulation [66,67].

In the herein presented computational model, the compared recombination strategies were assumed to be conferred by a single recombination modifier allele. Given the complexity of the recombination control system, this assumption is an apparent simplification even for constant recombination, let alone shift-inducible one. Yet, we still find such a simplification to be justified in a theoretical ‘proof-the-principle’ study. Our results suggest that a reversible plastic increase in RR in response to environmental perturbations can be beneficial under a wide range of selection parameters, even when such an increase emerges as a rough response to an external cue and does not account for finer genotype-specific differences. One of the crucial questions feeding the skepticism regarding the evolutionary importance of epigenetic factors is whether they are persistent enough to cause evolutionary effects [68]. The ever-increasing volume of empiric evidence on epigenetic variation includes numerous cases of long-term non-genetic transmission of epigenetic phenotypes. Much less clear is how frequently these effects can be accompanied by changes on the DNA level. An interesting fact is that the main factors participating in stress-induced epigenetic variation (microRNAs, chromatin changes, activation/silencing of transposable elements) are known players in the variation of genetic recombination, hence in transforming and more effective utilization of the population standing genetic variation. Yet, recombination is surprisingly rarely mentioned in the current rich literature on stress-induced inter- and trans-generational epigenetic variation. The herein presented results on shift-inducible recombination suggest that epigenetic factors may play an important role in the evolution of recombination plasticity. Our simulations demonstrate that it can evolve and affect population-level characteristics (such as mean recombination rate and mean fitness) even if the plastic changes lasted only one generation after shift.

## Acknowledgments

The authors thank the Hypernet Labs team for the access to the Galileo remote-computations platform and the kind support. SR is deeply thankful to Nataliya and Sandra Rybnikovs for their help in surveying the literature.

## Funding

The study was supported by the Israel Science Foundation (grant #1844/17 for AK, Graduate Studies Authority of the University of Haifa (SR), and the Israeli Ministry of Aliyah and Integration (SR).

## Notes

### Competing Interest Statement

The authors have declared no competing interest.

